# Alignment and quantification of ChIP-exo crosslinking patterns reveal the spatial organization of protein-DNA complexes

**DOI:** 10.1101/868604

**Authors:** Naomi Yamada, Matthew J. Rossi, Nina Farrell, B. Franklin Pugh, Shaun Mahony

## Abstract

The ChIP-exo assay precisely delineates protein-DNA crosslinking patterns by combining chromatin immunoprecipitation with 5′ to 3′ exonuclease digestion. Within a regulatory complex, the physical distance of a regulatory protein to DNA affects crosslinking efficiencies. Therefore, the spatial organization of a protein-DNA complex could potentially be inferred by analyzing how crosslinking signatures vary between the subunits of a regulatory complex. Here, we present a computational framework that aligns ChIP-exo crosslinking patterns from multiple proteins across a set of coordinately bound regulatory regions, and which detects and quantifies protein-DNA crosslinking events within the aligned profiles. By producing consistent measurements of protein-DNA crosslinking strengths across multiple proteins, our approach enables characterization of relative spatial organization within a regulatory complex. We demonstrate that our approach can recover aspects of regulatory complex spatial organization when applied to collections of ChIP-exo data that profile regulatory machinery at yeast ribosomal protein genes and yeast tRNA genes. We also demonstrate the ability to quantify changes in protein-DNA complex organization across conditions by applying our approach to data profiling *Drosophila* Pol II transcriptional components. Our results suggest that principled analyses of ChIP-exo crosslinking patterns enable inference of spatial organization within protein-DNA complexes.

## Introduction

Each cell type is defined by a unique gene expression program, which is in turn determined by the activities of regulatory proteins binding to promoters, enhancers, and other genomic regions. Genomic regulatory regions are bound by particular combinations of sequence-specific transcription factors (TFs), co-regulators, and chromatin modifiers in a spatiotemporal dependent manner. While large-scale efforts are underway to map and functionally characterize potential regulatory regions (Dunham *et al.*, 2012; Roadmap Epigenomics Consortium *et al.*, 2015), we still know relatively little about the structure and organization of individual protein-DNA complexes along the genome. To fully understand how gene regulatory programs are coordinated, it will be crucial to characterize precisely how regulatory complexes are assembled and organized.

Chromatin immunoprecipitation followed by sequencing (ChIP-seq) enables genome-wide localization of regulatory proteins. However, the spatial resolution of ChIP-seq is limited, as chromatin fragmentation strategies can result in sequencing reads that map several hundred base pairs away from the site bound by the protein of interest. Therefore, while integrative analyses of ChIP-seq data collections can find groups of co-bound regulatory proteins (Guo and Gifford, 2017; Giannopoulou and Elemento, 2013; Xie *et al.*, 2013), such analyses provide only limited insight into the spatial organization of proteins within regulatory complexes.

In contrast to ChIP-seq, ChIP-exo and related assays (e.g. ChIP-nexus (He *et al.*, 2015)), precisely define protein-DNA binding locations via the use of lambda exonuclease (Rhee and Pugh, 2011). The exonuclease digests protein-bound DNA in a 5′ to 3′ direction and, on average, stops at 6 bp before a protein-DNA crosslinking point. The ChIP-exo tag distribution at a given regulatory region is thus the product of crosslinking events that formaldehyde or other crosslinking agents have induced between the targeted protein and DNA.

ChIP-exo’s ability to map crosslinking signatures suggests a strategy for characterizing the spatial organization of regulatory complexes. At sites where a sequence-specific TF is bound directly to its cognate motif, the dominant crosslinking signature should result from direct interactions between the TF’s residues and proximal DNA bases. However, regulatory proteins that alternatively (or additionally) interact with DNA via protein-protein interactions should display crosslinking signatures related to the TFs that recruit them. Since the physical distance of a regulatory protein to the recruiting TF will affect crosslinking efficiencies, different members of a regulatory complex should display distinct crosslinking patterns across a regulatory region. In principle, then, analysis of ChIP-exo crosslinking patterns should enable some degree of inference regarding the spatial organization of regulatory proteins within a protein-DNA complex.

Previous work suggests that inferring the spatial organization of protein-DNA complexes via ChIP-exo crosslinking analysis is feasible. ChIP-exo analysis of yeast general transcription factors found that crosslinking patterns at Pol II promoters were consistent with those expected from crystallographic models of the transcriptional machinery (Rhee and Pugh, 2012). ChIP-exo characterization of ribosomal protein gene (RPG)-specific factors introduced the idea that the ordering of indirect protein-DNA interactions can be inferred from analysis of crosslinking efficiencies (Reja *et al.*, 2015). Specifically, high-resolution analysis of ChIP-exo crosslinking patterns at sites bound by the sequence-specific TF Rap1 show the same crosslinking pattern echoed in ChIP-exo experiments targeting Sfp1, Ifh1, and Fhl1, suggesting that these factors may be indirectly recruited to DNA by Rap1. We and others have made use of this concept to characterize indirect protein-DNA interactions in other systems. For example, in some mammalian cell types, both Glucocorticoid Receptor and Estrogen Receptor alpha may be indirectly recruited to certain binding sites via protein-protein interaction with FoxA TFs, as evidenced by near identical crosslinking patterns at those sites (Starick *et al.*, 2015; Yamada *et al.*, 2019). One limitation of previous approaches is that they have relied on TF binding motifs or known genomic anchor points to align bound sites before characterizing crosslinking patterns. (Rhee and Pugh, 2011; Rossi *et al.*, 2018; Reja *et al.*, 2015). Naturally, such strategies limit the usefulness of ChIP-exo crosslinking analysis to protein-DNA complexes where the focal point of spatial organization is already known.

In this work, we formalize concepts suggested by previous studies by presenting ChExAlign, a systematic approach for characterizing ChIP-exo crosslinking patterns across multiple members of a protein-DNA regulatory complex. Specifically, we first develop a multiple alignment procedure for characterizing consistent ChIP-exo crosslinking signatures across multiple ChIP-exo experiments and across multiple regulatory regions. To make the alignment approach broadly applicable to different types of protein-DNA complexes, our procedure does not rely on sequence features or other genomic annotations, but rather directly aligns multi-protein ChIP-exo tag profiles. While there has been limited work on aligning broad tag distributions such as those from histone modification ChIP-seq and ChIP-chip data (Lai and Buck, 2010; Hon *et al.*, 2008; Nielsen *et al.*, 2012; Nair *et al.*, 2014), our approach is optimized for high-resolution ChIP-exo data. We use an overlap Needleman-Wunsch alignment (Needleman and Wunsch, 1970; Durbin *et al.*, 2010) to progressively align per-base, strand separated ChIP-exo tag profiles. Given a multiple alignment of ChIP-exo tag profiles, our approach next applies a probabilistic mixture model to deconvolve individual protein-DNA crosslinking events. This approach allows consistent quantification of crosslinking strengths across multiple proteins in a regulatory complex. Finally, we apply principal component analysis (PCA) to visualize similarities between the crosslinking preferences of the regulatory proteins.

We demonstrate the utility of our approach by applying it to characterize the spatial organization of three distinct regulatory complexes. We first apply our method to RPG ChIP-exo datasets in order to show that multiple profile alignment can be used to automatically align collections of ChIP-exo data across a collection of coordinately regulated regions. While previous analyses of the RPG ChIP-exo data relied on a manual alignment around a sequence motif feature (Rap1 binding sites), our ChIP-exo alignment approach yields similar alignments, and the same biological conclusions, without knowledge of DNA sequence features. Secondly, we extend our analyses to 12 novel ChIP-exo datasets that characterize the occupancy of regulatory complexes at tRNA genes. Due to the variable length of tRNA intragenic promoters, we extend our multiple profile alignment procedure to account for affine gaps. We further demonstrate that crosslinking event detection and quantification yields insight into the spatial organization of individual proteins within the Pol III transcriptional machinery. Finally, we demonstrate that crosslinking analysis provides a quantitative framework for characterizing changes in regulatory complex organization across conditions. By applying our methods to a collection of ChIP-nexus data that profile *Drosophila* Pol II transcriptional components under two experimental conditions, we demonstrate that we can quantify the degree to which individual proteins are relocalized when transcriptional initiation is inhibited.

In summary, our approaches provide a novel platform for examining the spatial organization of protein-DNA complexes from collections of high-resolution ChIP data.

## Results

### ChIP-exo profile alignment recovers the spatial organization of a regulatory complex at ribosomal protein genes

In order to demonstrate that ChExAlign provides an informative alignment of protein-DNA crosslinking patterns across sets of related regulatory regions, we first applied it to analyze the organization of a protein-DNA complex at yeast ribosomal protein genes (RPGs). Most yeast RPGs are coordinately regulated by a common set of regulatory proteins. The transcription factor Rap1 binds to cognate sequence motifs located 77bp-501bp upstream of the transcription start site (TSS), and recruits Fhl1, Ifh1, and Sfp1. Hmo1 is also recruited at roughly half of the RPGs (Knight *et al.*, 2014; Reja *et al.*, 2015). Previous analyses of Rap1, Flh1, Ifh1, Sfp1, and Hmo1 ChIP-exo data determined that the RPG regulatory complex has a well-defined spatial organization (Reja *et al.*, 2015). Rap1 binding sites serve as an upstream boundary to the complex. Fhl1, Ifh1, and Sfp1 are almost identically positioned ~100bp downstream of Rap1, with some evidence of additional crosslinking through the Rap1 site (indicative of protein-protein interactions between Rap1 and the recruited factors). When present, Hmo1 occupies the region between Rap1 and the TSS, essentially overlapping where Fhl1/Ifh1/Sfp1 bind. Thus, a consistently organized regulatory complex is present in the upstream regions of most yeast RPGs, and this organization should in principle be recoverable by aligning ChIP-exo profiles that span the RPG regulatory regions.

A standard approach to analyzing collections of regulatory genomics data at a set of related gene loci might begin by producing composite profiles centered on the genes’ TSSs. Applying this approach to the five ChIP-exo datasets at 134 RPGs produces a set of smooth composite profiles without any discernable organization between the members of the regulatory complex (Fig 1A). Indeed, previous analysis has demonstrated that it is only when the five ChIP-exo datasets are aligned by Rap1-bound motif locations (consistently oriented with respect to the RPG TSSs) that a more structured organization emerges from the data (Reja *et al.*, 2015). Thus, the smooth profiles produced by a TSS-centric alignment are artefacts of the variable spacing between TSSs and the true organizing points of the regulatory complex (i.e. the Rap1 sites). We therefore asked whether ChExAlign’s alignment procedure can recapitulate insights into the organization of the RPG regulatory complex without using sequence motif information or prior knowledge of Rap1 sites as the organizing loci.

**Figure 1.**
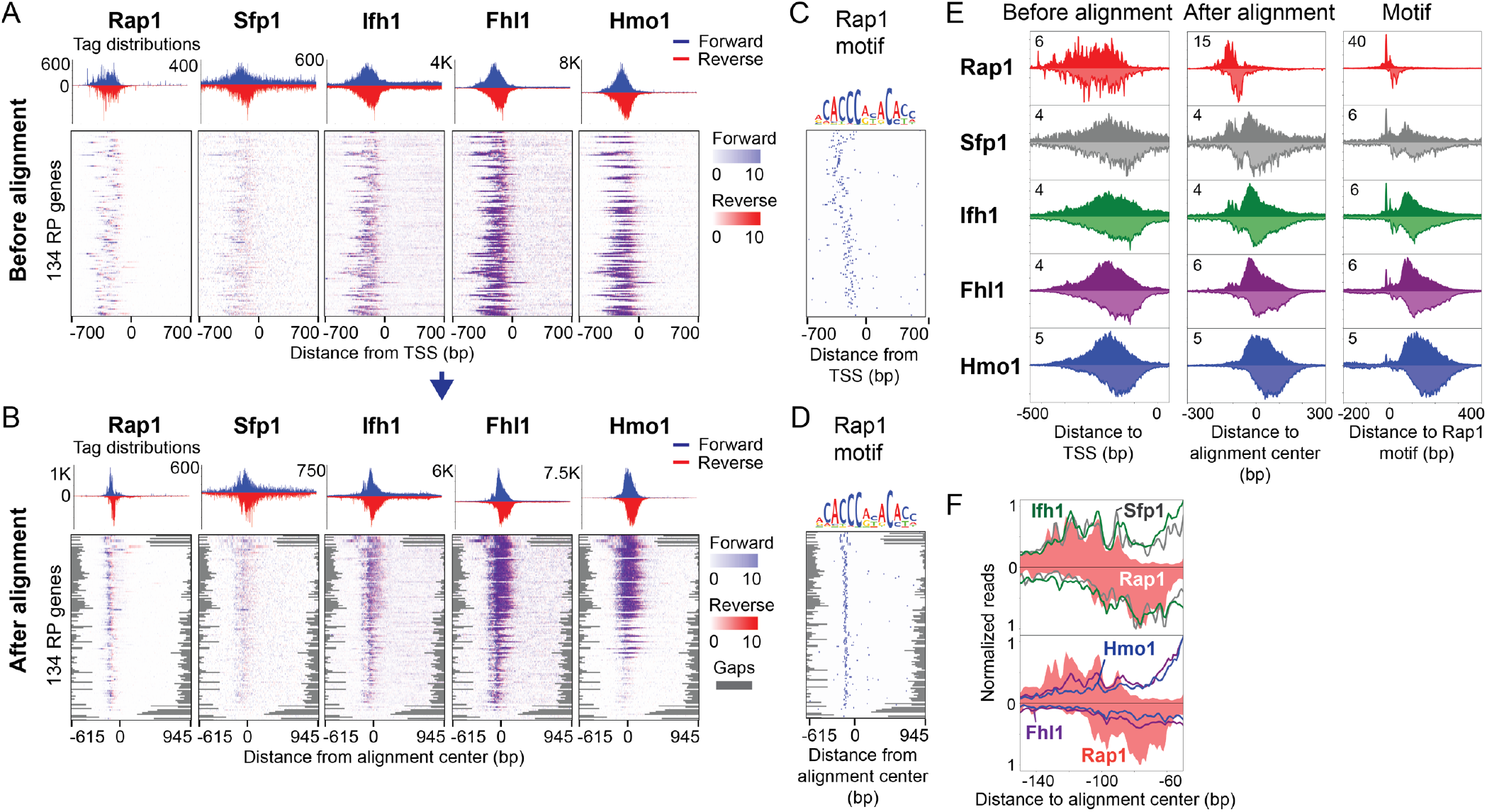
ChIP-exo tag alignment and an inferred organization of RPG specific factors. **A) B)** Rap1, Sfp1, Ifh1, Fhl1, and Hmo1 ChIP-exo enrichment at 134 RP genes, centered on TSS coordinates (A) and after ChExAlign alignment (B). Forward strand tags are shown in blue and reverse strand tags are shown in red. The heatmaps are sorted based on the order by which alignment was performed (B). **C)** Positions of Rap1 motif in the same windows displayed in A). **D)** Positions of Rap1 motif given the aligned coordinates displayed in B). **E)** ChIP-exo tag patterns of RPG-specific factors before and after alignment, and centered around Rap1 motif. **F)** ChIP-exo tag patterns for Sfp1, Ifh1, Fhl1, Hmo1 (grey, green, purple, and blue traces) compared with the equivalent of Rap1 (pink-filled plots) after progressive alignment.

We applied ChExAlign to produce an ungapped overlap multiple profile alignment of 5-dimensional ChIP-exo tag profiles (Rap1, Flh1, Ifh1, Sfp1, and Hmo1) taken from 1,400bp windows centered on the TSSs of 134 yeast RPGs (Fig. 1B). Our alignment recovers sharply distributed composite profiles across the five factors (Fig. 1B, 1E). The aligned composite plots enable some degree of inference regarding the organization of the regulatory complex. For example, high-resolution analysis of the crosslinking patterns displayed at the Rap1 binding sites suggest that Fhl1/Ifh1/Sfp1 are indirectly bound through protein-protein interactions with Rap1 (Fig. 1F). In addition, the multiple profile alignment appropriately clusters the subset of RPGs that displays Hmo1 enrichment, consistent with previous observations (Knight *et al.*, 2014; Reja *et al.*, 2015). Importantly, even though ChExAlign does not use sequence information during the alignment procedure, the multiple profile alignment induces an alignment of the underlying Rap1 motif sequences (Fig. 1C, 1D). Therefore, ChExAlign can accurately align multi-dimensional ChIP-exo profiles across a set of coordinately regulated regions, enabling insights about the organization of regulatory proteins within the regions.

### Gapped ChIP-exo profile alignment enables consistent analysis of protein-DNA crosslinking patterns across 12 regulatory proteins at tRNA genes

Having demonstrated that multiple profile alignment can recover informative protein-DNA crosslinking patterns in a set of regions that share a tightly organized regulatory complex, we next aimed to demonstrate that the alignment procedure is robust to cases where the protein-DNA regulatory complex contains a more variably spaced organization. We chose to focus on protein-DNA interactions in the yeast Pol III transcriptional machinery of tRNA genes, as tRNA genes with varying lengths contain regulatory elements of consistent composition but variable internal spacing. Analyses of protein-DNA crosslinking patterns over tRNA might therefore be expected to require gapped ChIP-exo profile alignment strategies.

Eukaryotic tRNA genes are transcribed by Pol III, which is recruited by the multi-subunit transcription factor complexes TFIIIB and TFIIIC (Deprez *et al.*, 1999; Kassavetis *et al.*, 1990) (Fig 2A). TFIIIB is composed of TBP, Brf1, and Bdp1, while TFIIIC contains two subcomplexes, τA (composed of Tfc1, Tfc4, and Tfc7) and τB (composed of Tfc3, Tfc6, and Tfc8). The TFIIIC subcomplexes bind to conserved intragenic promoter motifs, named Box A (bound by τA) and Box B (bound by τB), and enable assembly of the TFIIIB complex at a region approximately 30bp upstream of the tRNA transcription start site (Dumay-Odelot *et al.*, 2002; Rüth *et al.*, 1996; Chaussivert *et al.*, 1995). Pol III is then recruited by TFIIIB, enabling transcription. Yeast tRNA genes vary in length between 74bp and 134bp, and this variation is reflected by variable spacing between the intragenic Box A and Box B promoter elements (Fig 2B).

**Figure 2.**
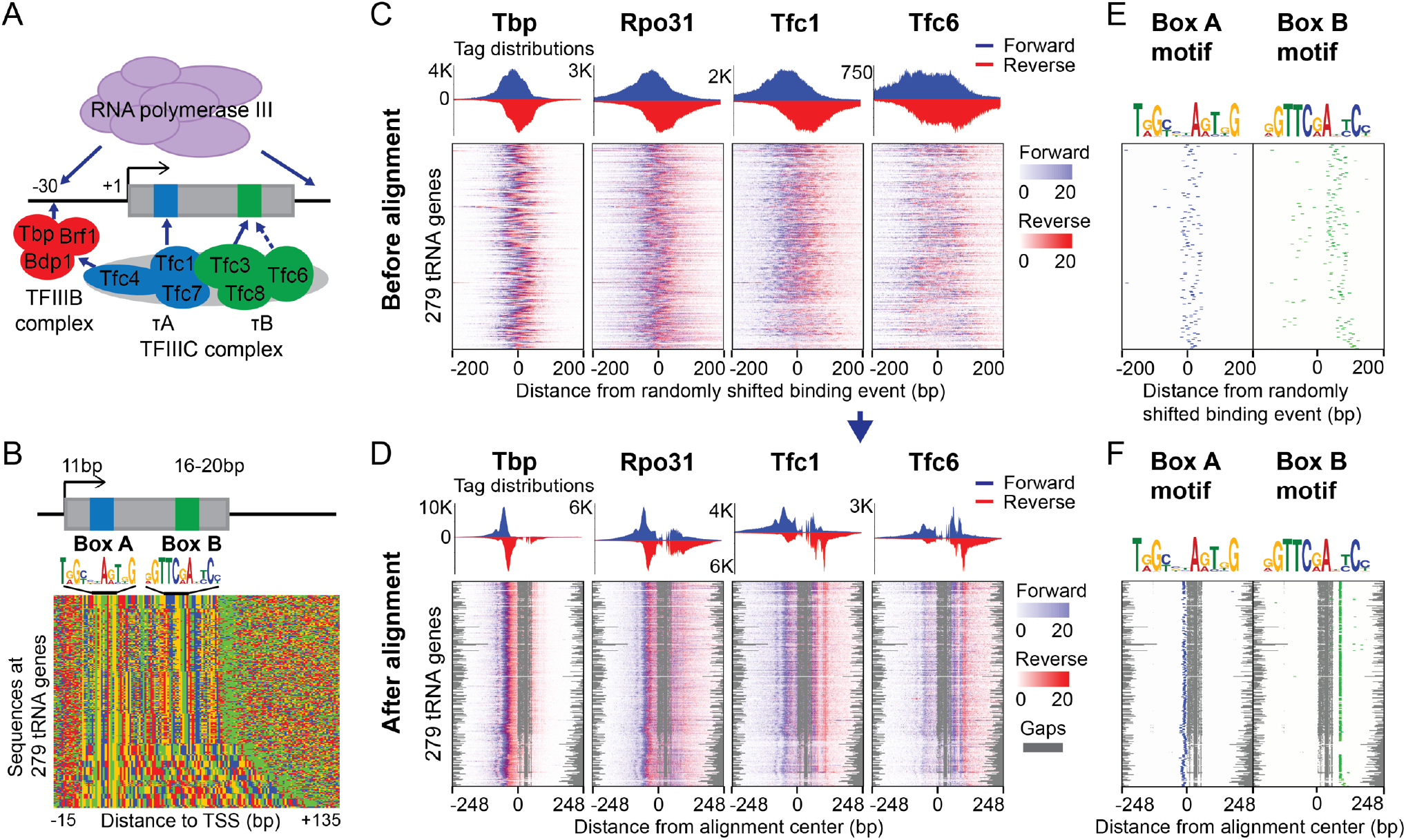
Gapped alignment of ChIP-exo profiles enables characterization of consistent protein-DNA crosslinking patterns across 279 yeast tRNA genes. **A)** Cartoon representation of protein organization within the yeast tRNA transcriptional machinery. **B)** Sequence plot of tRNA genes, sorted by increasing tRNA gene length. The TSS and Box A are separated by 11bp, while Box B and the transcription end site are separated by 16-20bp. **C), D)** The alignment method simultaneously aligns 12 ChIP-exo experiments with gaps. ChIP-exo heatmaps before (C) and after (D) alignment are shown for TBP (TFIIIB), Rpo31 (Pol III), Tfc1 (TFIIIC τA), and Tfc6 (TFIIIC τB). Gaps are represented in grey. The heatmaps are sorted by increasing tRNA gene length. **E), F)** Relative positions of Box A and Box B motifs before (E) and after (F) the alignment.

We applied ChIP-exo to generate novel high-resolution protein-DNA interaction profiles for three protein components of Pol III, the three components of TFIIIB, and the six components of TFIIIC (Fig 2C, 2D). In order to account for the expected variable spacing between crosslinking signatures centered on the Box A and Box B sites in tRNA genes of different lengths, we extended our ChIP-exo multiple profile alignment strategy to incorporate affine gap penalties. We then applied our framework to align ChIP-exo profiles from the 12 ChIP-exo targets across 279 yeast tRNA gene loci. Our goal is to demonstrate that our ChIP-exo profile alignment procedure can recover informative protein-DNA crosslinking signatures for a regulatory complex without any prior knowledge of sequence features. However, the tRNA genes contain highly conserved sequence features, so initializing the alignments centered on tRNA TSSs would run the risk of being indirectly biased by the underlying conserved sequences (Fig 2B). We therefore chose to remove this potential confounding effect by initializing the alignments as centered on randomly shifted locations within a 60bp range surrounding tRNA TSSs (Fig. 2C). This also has the effect of scrambling the relative locations of the Box A and Box B motifs in the initial alignment (Fig. 2E).

The aligned ChIP-exo profiles produced by ChExAlign multiple profile alignment display pronounced peaks compared to the initial alignment (Fig. 2C, 2D; representative profiles for protein components of Pol III, TFIIIB, and TFIIIC τA & τB are shown for illustration). Despite the fact that no sequence information is included in the alignment procedure, the multiple profile alignment induces a tight alignment of both Box A and Box B motif locations (Fig. 2F). The aligned ChIP-exo profiles incorporate a gap between the Box A and Box B motifs; as expected, short tRNA genes contain a larger gap compared with longer tRNA genes, having the effect that a consistent crosslinking profile alignment is constructed across all tRNA gene loci (Fig 2D, 2F). Visual inspection shows that the multiple profile alignment contains sharp protein-DNA crosslinking peaks near the major regulatory elements (i.e. surrounding the Box A motif, Box B motif, and upstream TFIIIB binding location), consistent with where formaldehyde-induced protein-DNA crosslinking events would be expected to occur for the profiled regulatory proteins. Our results therefore demonstrate that ChExAlign can produce a high quality multiple profile alignment of ChIP-exo crosslinking signatures, even in cases where the underlying regulatory elements occur at variable spacing in the constituent genomic loci.

### ChIP-exo crosslinking quantification enables inference of protein-DNA complex organization

Visual inspection of aligned ChIP-exo crosslinking profiles enables some degree of insight into the spatial organization of protein-DNA complexes. For example, the aligned crosslinking profiles of tRNA transcriptional components show that TBP and other TFIIIB components are primarily crosslinked through a site upstream of the TSS, as expected from the direct protein-DNA crosslinks resulting from TBP binding to the TATA motif (Fig. 3A). Meanwhile, the TFIIIC components primarily display crosslinking through sites around the Box A and Box B motifs, again reflective of the expected protein-DNA binding events. However, TFIIIC components also display additional, weaker crosslinking signatures through the same site that is crosslinked by TFIIIB components, indicative of the indirect protein-DNA crosslinking that results from protein-protein interactions between TFIIIC and TFIIIB. Within a regulatory complex, we should expect a protein’s crosslinking efficiency within a given DNA site to decay as a function of the number of protein-protein crosslinks required to link it to the protein that directly contacts the DNA site. Therefore, careful analysis of the relative strengths of all crosslinking peaks may enable inference of protein spatial positioning within a regulatory complex. To facilitate such inference, we aimed to incorporate a principled approach to quantifying and visualizing crosslinking peaks into the ChExAlign framework.

Our approach to quantifying ChIP-exo crosslinking signatures first applies a probabilistic mixture model to estimate the positions and relative strengths of crosslinking events within a multiple ChIP-exo profile alignment (see Methods). The mixture model probabilistically assigns observed ChIP-exo tags to crosslinking events using a predefined model of tag distributions around single crosslinking events. We use the Expectation Maximization (EM) algorithm to iteratively optimize the positions and strengths of crosslinking events using information from the assigned tags. The mixture model incorporates priors to keep the number of detected crosslinking events low (i.e. assuming that protein-DNA crosslinking occurs at relatively few bases in a regulatory region) and to keep the positions of crosslinking events consistent across the aligned multi-protein profiles (i.e. assuming that the same crosslinking events will recur across ChIP-exo profiles from multiple interacting proteins in a regulatory complex).

We applied our crosslinking deconvolution procedure to the 12-dimensional multiple profile alignment of the tRNA regulatory complex (Fig. 3A), first subtracting scaled ChIP-exo control signals from the individual profiles to account for elevated ChIP-exo background signals over the tRNA genes. The result of the deconvolution procedure is illustrated for Tfc6 in Fig. 3B; crosslinking events are detected at 8 positions within the Tfc6 signature, and each has a relative strength quantification which is derived from the proportion of Tfc6 ChIP-exo tags that is associated by the mixture model. The results of the procedure across all profiles can be summarized using a relative crosslinking matrix (Fig. 3C). The matrix illustrates that TBP has its highest crosslinking signal at position −60 in the alignment, a position that is consistent with TBP’s cognate binding motif (Fig. 3C). Brf1 and Pol III components also show the strongest signal at the same position, suggesting that they associate with DNA via protein-protein interactions with TBP (Fig. 3C). On the other hand, Tfc6 and Tfc8 from the TFIIIC τB subcomplex show highest signals at positions +47 and +77, again consistent with their role in binding the Box B motif.

**Figure 3.**
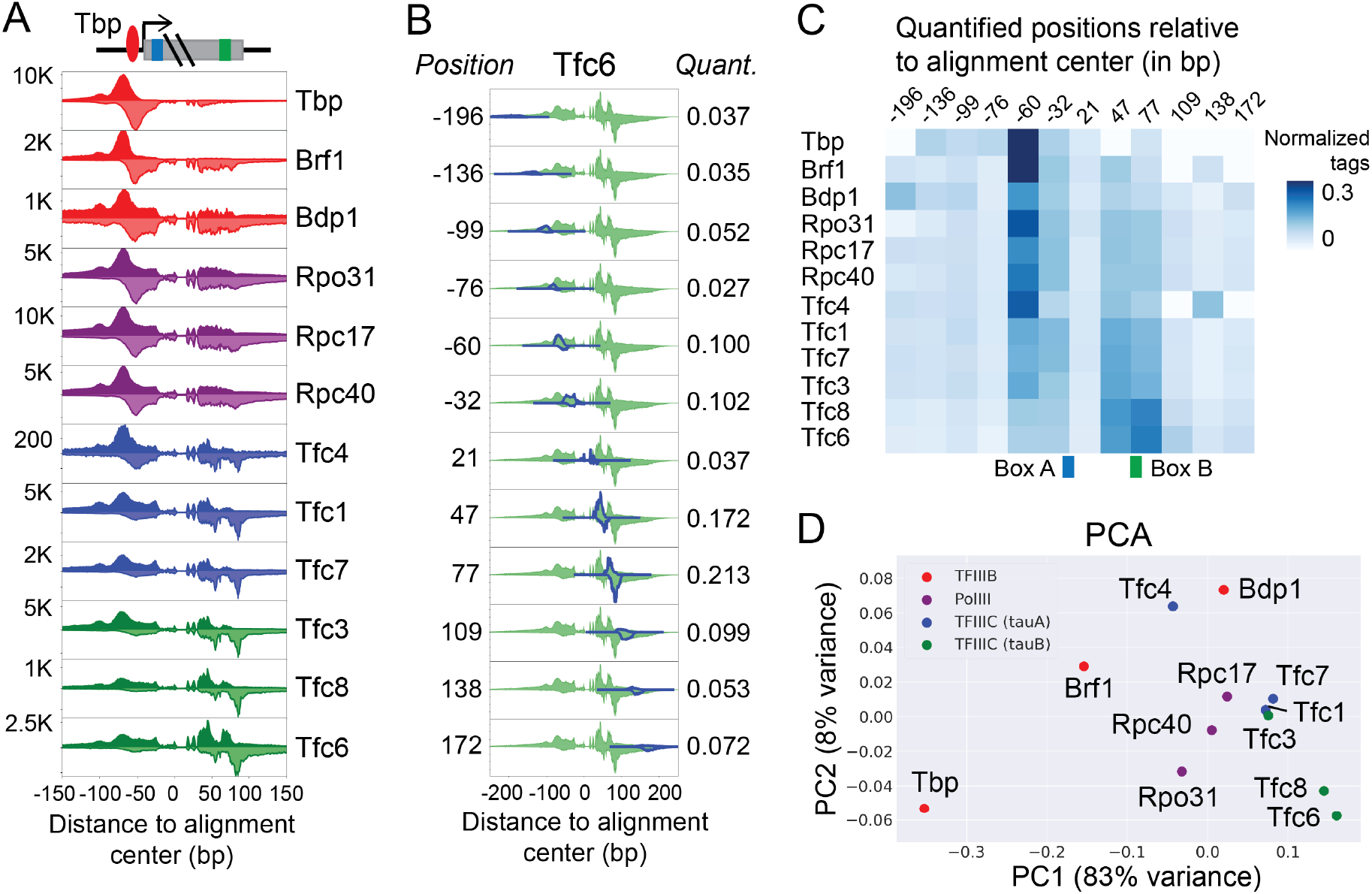
Inferring the positional organization of the tRNA transcriptional multi-subunit complex using ChIP-exo crosslinking quantification. **A)** Aligned ChIP-exo profiles of Pol III, TFIIIB, and TFIIIC components at tRNA genes show distinct crosslinking patterns. **B)** ChExAlign uses a mixture model to deconvolve the positions and relative strengths of protein-DNA crosslinking events across the aligned multi-protein ChIP-exo profiles. The effect of crosslinking event deconvolution in the Tfc6 ChIP-exo profile is shown as an example. **C)** Matrix of relative crosslinking strengths for TFIIIB, Polymerase III, and TFIIIC components at detected crosslinking event positions. **D)** Principal Component Analysis applied to the matrix of relative crosslinking strengths approximates aspects of the known organization of the yeast tRNA transcriptional machinery (Fig. 2A). PC1 and PC2 together explain 91% of the variance.

To visually summarize all crosslinking quantifications, we applied Principal Component Analysis (PCA) to the rows of the crosslinking matrix (Fig. 3D). This dimensionality reduction organizes the proteins in a manner that resembles the expected spatial organization of the tRNA transcriptional complex (Fig. 2A). TFIIIB components TBP and Brf1 are the left-most outliers of the plot, while Pol III and TFIIIC complexes are grouped in the middle and the right side of the plot. Notably, Tfc4, which is known to crosslink to both TFIIIB and to the rest of the τA subcomplex (Liao *et al.*, 2003, 2006; Male *et al.*, 2015; Chaussivert *et al.*, 1995) is situated between Brf1 and other τA subcomplex members on the PCA plot. Consistent with a previous study that showed Bdp1 crosslinking to the C34 subunit of Pol III (Khoo *et al.*, 2014), Bdp1 is situated proximal to Pol III subunits on the PCA plot. Moreover, in accordance with the finding that Tfc3 crosslinks with Tfc1 and Box B (Male *et al.*, 2015), the PCA plot shows Tfc3 in between τA and τB subunits of TFIIIC.

In summary, ChExAlign’s crosslinking deconvolution and quantification procedures, and downstream dimensionality reduction, are promising approaches for generating hypotheses regarding the spatial organization of a protein-DNA complex from a collection of ChIP-exo data.

### ChExAlign enables a principled quantification of regulatory complex reorganization across conditions

If the structure of a regulatory complex changes across different gene classes or between cell types, we should expect that the relative crosslinking signatures of members of the complex will also change. Provided that data from all conditions are analyzed simultaneously, ChExAlign’s approach to aligning and quantifying protein-DNA crosslinking signatures is suitable for detecting spatial localization changes within a regulatory complex across conditions. To demonstrate our framework’s ability to detect regulatory complex reorganization, we applied it to ChIP-nexus data profiling Pol II, the basal transcription factors TBP, TFIIA, TFIIB, TFIIF, TAF2, XPB, and negative elongation factor E (NELFE) in *Drosophila melanogaster* Kc167 cells (Shao and Zeitlinger, 2017). Shao & Zeitlinger collected data profiling these factors both under control conditions and after applying Triptolide (TRI) treatment, which blocks transcriptional initiation. We therefore sought to apply ChExAlign to this collection of ChIP-nexus datasets to quantify TRI-dependent changes in protein-DNA crosslinking profiles across factors.

We applied ChExAlign to align 16 ChIP-nexus profiles (eight factors, each in two conditions) across the top 500 most enriched Pol II peak locations (Fig. 4A, 4C). The multiple profile alignment procedure tightly aligns ChIP-nexus profiles across these factors and conditions, as indicated by the high consistency between our alignment and annotated gene TSSs in the regions; 99% regions were aligned in the same orientation as the annotated TSS, and 53% of annotated TSSs were within 10bp of what we defined to be the zero position of the multiple profile alignment (Fig. 4B). We next used our mixture model approach to quantify crosslinking strengths for each factor and across conditions (e.g. Fig. 4D).

**Figure 4.**
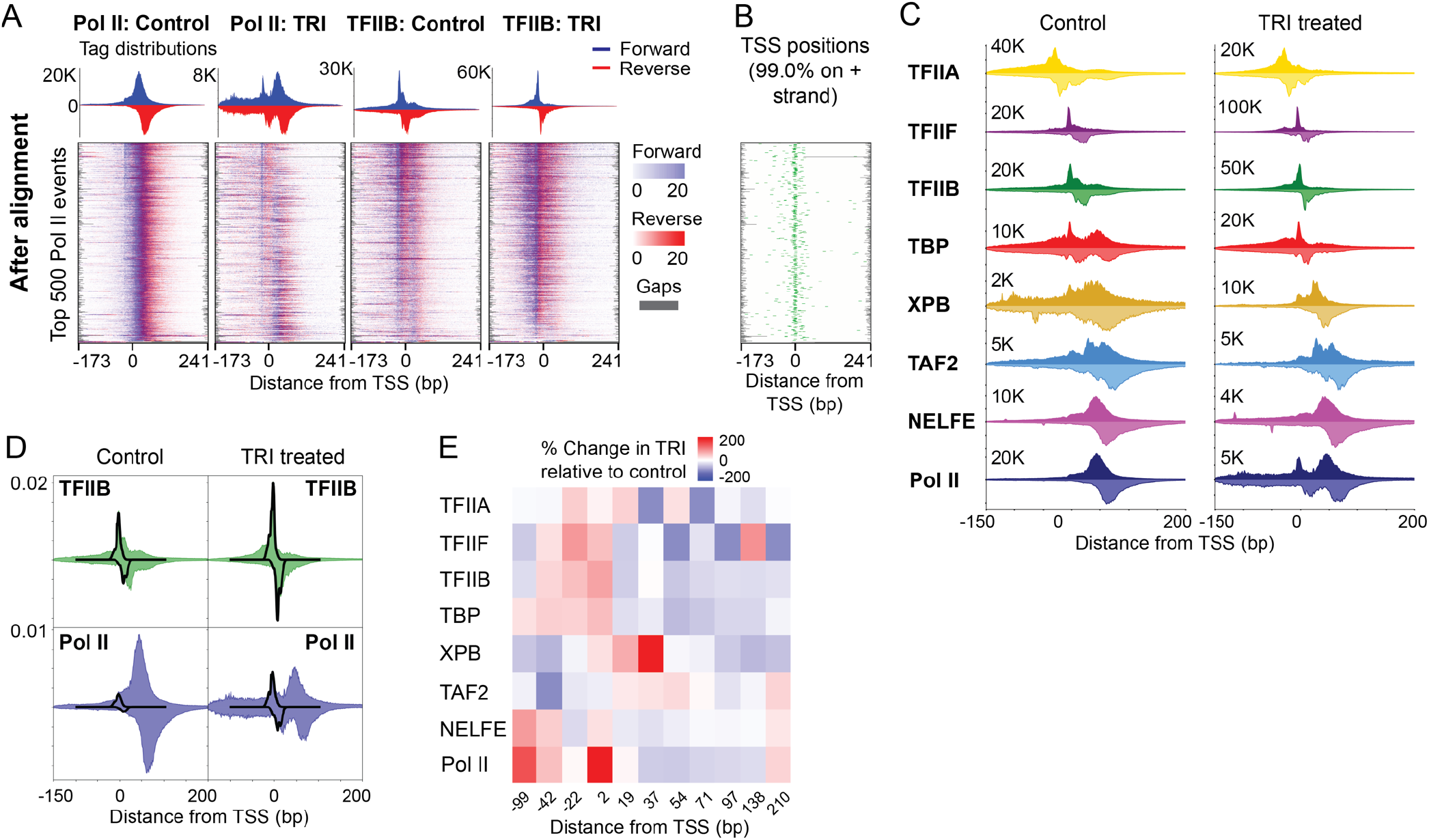
ChExAlign enables quantification of regulatory complex reorganization across protein components and conditions. **A)** ChIP-nexus multiple profile alignments for Pol II and TFIIB in control and TRI treatment conditions. Alignments are performed across the top 500 Pol II binding event locations. Heatmaps are sorted based on the order resulting from the multiple alignment. **B)** Annotated *Drosophila* gene TSS positions plotted relative to the multiple profile alignment. **C)** Aligned ChIP-nexus tag patterns of eight factors with or without TRI treatment. **D)** Example of ChIP-nexus tag deconvolution at the position +2 over the TFIIB and Pol II profile (control and TRI treatment condition). **E)** Percent difference in crosslinking strengths for eight factors at detected crosslinking event positions. Red represents increase in crosslinking strength for TRI treatment, while blue shows increase in crosslinking for control condition.

As illustrated in Fig. 4C, Pol II’s crosslinking profile in the control condition shows one major peak at alignment position +54, while the profile in the TRI treatment condition shows additional crosslinking at alignment position +2. This apparent shift of Pol II occupancy is explained in detail in the source publication; the shift in crosslinking to the +2 position is due to the accumulation of Pol II at the pre-initiation complex (PIC) when it is prevented from moving into the downstream transcriptional pause site by TRI treatment (Shao and Zeitlinger, 2017). Similarly, an increase in TFIIB occupancy is observed at position +2 in the TRI treatment, condition compared to the control. Our approach allows us to quantify this effect. Comparison of crosslinking quantification across conditions shows a 2-fold increase in Pol II crosslinking strength at the +2 PIC position after TRI treatment, while the +54 pause site position shows a 47% decrease in crosslinking strength. Other changes discussed in the original publication are also provided with a quantitative basis by our approach. For example, XPB, the direct target of TRI treatment, shows a 2-fold increase in crosslinking strength at the +37 position after TRI treatment, consistent with it blocking transcriptional initiation at that site. Similarly, a 60% decrease in TBP crosslinking at the +54 position after TRI treatment is consistent with TBP enrichment at that location being dependent on Pol II moving into the pause site. While these factors show distinct differences between the conditions, factors such as NELFE and TAF2 retained similar crosslinking patterns after the TRI treatment (Fig 4E). Our results thus demonstrate that crosslinking profile alignment followed by mixture model analysis enables us to quantify changes in regulatory complex organization between different conditions as well as across different experimental targets.

## Discussion

We have presented ChExAlign, a principled framework for characterizing ChIP-exo crosslinking patterns across multiple members of a protein-DNA complex. Our approach implements a multiple alignment procedure that is optimized for aligning multi-dimensional ChIP-exo profiles, and which does not rely on sequence motif or genomic annotation information. ChExAlign also encapsulates a novel mixture modeling approach for estimating the locations and relative strengths of crosslinking events within a set of aligned ChIP-exo profiles. As we have demonstrated, the resulting crosslinking matrix can be used to visualize the relative spatial relationships between members of a protein-DNA complex (via dimensionality reduction), and to quantify shifts in spatial positioning across experiment conditions.

By applying ChExAlign to characterize the spatial organization of three distinct regulatory complexes, we have demonstrated that our approach is generally applicable to collections of ChIP-exo data profiling multiple members of a protein-DNA complex that is consistently organized across a set of genomic regions. Several previous studies have proposed the idea of characterizing the organization of protein-DNA complexes from ChIP-exo data (Reja *et al.*, 2015; Shao and Zeitlinger, 2017; Rhee *et al.*, 2016). However, these previous approaches were typically limited to aligning data manually around known focal points of the protein-DNA complex (e.g. TF binding motifs or genomic annotations) and thus become impractical when analyzing large protein-DNA complexes that may be crosslinked at multiple unknown locations within regulatory regions.

The gold standard for characterizing the spatial organization of regulatory complexes is the creation of 3-dimensional structural models by applying structural biology techniques such as X-ray crystallography or cryo-EM to purified complexes. Our approaches by no means result in the same type of structural information. However, careful analysis of protein-DNA crosslinking patterns might add useful orthogonal information to structural studies. For example, ChIP-exo crosslinking analysis might help to confirm the *in vivo* relevance of 3-dimensional structures that have been defined *in vitro*. Similarly, analysis of how crosslinking patterns vary across regulatory regions or experimental conditions may help to clarify how consistent the structure of a regulatory complex is in biological conditions.

In summary, we have demonstrated that our framework enables new forms of insight from analysis of multiple ChIP-exo crosslinking patterns, taking another step towards the fine-grained characterization of spatial organization within protein-DNA complexes. As larger collections of high-resolution protein-DNA interaction data become available, our analysis framework will contribute to investigating transcriptional mechanisms in greater detail.

## Methods

### Pairwise alignment of ChIP-exo profiles with affine gap score

To align multi-experiment ChIP-exo tag profiles across multiple genomic regions, we extend an affine gapped, overlap alignment version of the Needleman-Wunsch algorithm (Needleman and Wunsch, 1970; Durbin *et al.*, 2010). The Needleman-Wunsch algorithm is popularly used to perform global alignment of protein or nucleotide sequences. Here, rather than aligning discrete character sequences, our approach aligns real-valued matrices of ChIP-exo tag counts. For a given length *L* genomic region, the corresponding input matrix contains rows for each of the two DNA strands in each of *K* ChIP-exo experiments. The elements in each row contain the number of ChIP-exo tag 5′ positions (stranded) mapped to each base in the region. Thus, the dimensionality of each input matrix is 2*K* × *L*, and each ChIP-exo tag is counted in only a single bin of the matrix. The goal of the pairwise alignment procedure is to form a global alignment between ChIP-exo profile matrices corresponding to genomic regions *X* = (*x*_*1*_, …, *x*_*i*_, …, *x*_*L*_) and *Y* = (*y*_*1*_, …, *y*_*j*_, …, *y*_*L*_).

As in the original dynamic programming alignment with affine gap penalties, we construct matrices *M*, *I*_*x*_, and *I*_*y*_, indexed by *i* and *j*, where the value *M*(*i*, *j*) is the best score up to (*i*, *j*) given that *x*_*i*_ is aligned to *y*_*j*_, *I*_*x*_(*i*, *j*) is the best score given that *x*_*i*_ is aligned to a gap, and *I*_*y*_(*i*, *j*) is the best score given that *y*_*j*_ is an insertion with respect to *x*. We use an affine gap cost structure to impose different penalties for opening and extending a gap of length *g* as γ(*g*) = −*d* − (*g* − 1)*e*, where *d* is the *gap-open* penalty and *e* is the *gap-extension* penalty. In this study, we use *e* = 0.1*d*. We use *d*=50 in analyzing Pol III transcriptional complex at tRNA genes, *d*=100 for ChIP-nexus data, and *d*=200 for analyzing RPG-specific factors. The recursion relationships are unchanged from the original algorithm:

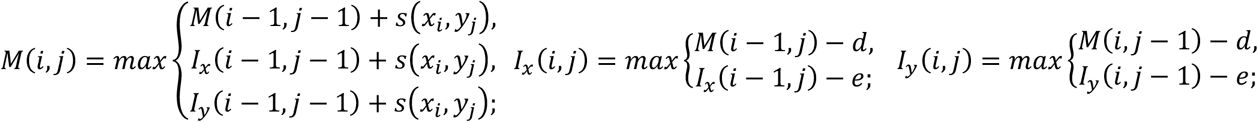

The similarity score of *x*_*i*_ and *y*_*j*_ coordinates, *s*(*x*_*i*_, *y*_*j*_) is computed using the Pearson correlation coefficient as follows:

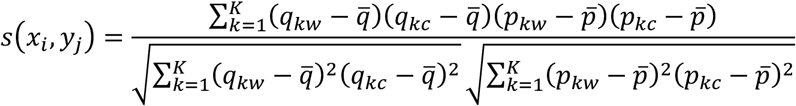

where 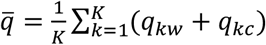 and 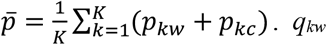. *q*_*kw*_ and *q*_*kc*_ are the numbers of ChIP-exo tag 5′ positions mapped from experiment *k* to the Watson and Crick strands (respectively) at the *x*_*i*_ coordinate. *p*_*kw*_ and *p*_*kc*_ are the numbers of ChIP-exo tag 5′ positions mapped from experiment *k* to the Watson and Crick strands (respectively) at the *y*_*j*_ coordinate.

Our approach aims to generate overlap alignments, which do not penalize overhanging ends. Therefore, the scoring matrices are initialized with *M*(*i*, 0)=0, *I*_*x*_(*i*, 0)=0, and *I*_*y*_(*i*, 0)=0 for *i*=1,…,*n* and *M*(0, *j*)=0, *I*_*x*_(0, *j*)=0, and *I*_*y*_(0, *j*)=0 for *j*=1,…,*m*. Traceback starts from the cell with the maximum value on the lower right quadrant border, (i.e. (*n*, *j*), *j*=2/*m*,…,*m* or (*i*, *m*), *i*=2/*n*,…,*n*), among the *M*, *I*_*x*_, and *I*_*y*_ matrices. Traceback continues until the top or left edge is reached.

### Progressive alignment of multiple regions

To build multiple alignments, we progressively align nodes by a succession of pairwise profile alignments. Nodes initially contain single region profiles. Progressive alignment proceeds by aligning the most similar pair of profiles, and merging the aligned pair into a composite profile generated according to the gapped alignment. Gas are represented by one dimensional array with size of alignment length. Similarities are then recalculated between the aligned profile and all remaining regions. We repeat these steps until all the nodes are merged. After the final multiple alignment of all regions has been constructed, we subtract background signals from the composite ChIP-exo profiles. We first use NCIS (Liang and Keles, 2012) to scale a control experiment to each analyzed signal ChIP-exo experiment. We then subtract scaled per-base control tag counts from each signal ChIP-exo experiment.

### Deconvolution and quantification of crosslinking profiles using a mixture model

After constructing composite ChIP-exo profiles representing the multiple alignment of analyzed regions, we aim to locate and quantify protein-DNA crosslinking positions within the profiles. Our approach models ChIP-exo composite data as being generated by a mixture of crosslinking events, and an Expectation Maximization (EM) learning scheme is used to probabilistically assign sequencing tags to crosslinking positions. By estimating crosslinking positions in the composite profiles, we implicitly assume that protein-DNA crosslinking patterns are consistent at all aligned regions, albeit with spacing differences between crosslinking sites as modeled by the alignment gaps.

Pr(*r*_*n*_|*x*) gives the probability of observing ChIP-exo tag *r*_*n*_ from a crosslinking event located at genomic coordinate *x* and is defined by a pair of Gaussian distributions (one on the positive strand, one on the negative strand) with σ = 6bp and offset from each other by 12bp (positive strand to the left). We define a vector of component locations ***µ*** where *µ*_*j*_ is the genomic location of the crosslinking event *j*. We initialize potential crosslinking events such that they are spaced in 5bp intervals along the window. The overall likelihood of the observed set of tags, ***r***, given the crosslinking event positions, ***µ***, the binding event mixture probabilities (i.e. crosslinking event strengths), ***π***, is given by:

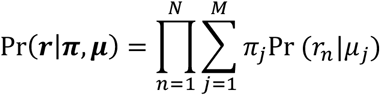

To limit the number of modeled crosslinking positions, we place a sparseness promoting negative Dirichlet prior, *α*, on the crosslinking strength ***π*** (Figueiredo and Jain, 2002). The latent assignments of tags to crosslinking events are represented by the vector ***z***. The complete-data log posterior is as follows:

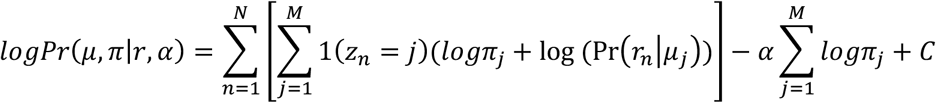

The E-step that calculates the relative responsibility of each crosslinking event in generating each tag is:

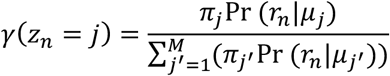

The maximum a posteriori probability (MAP) estimation (Figueiredo and Jain, 2002) of ***π*** is:

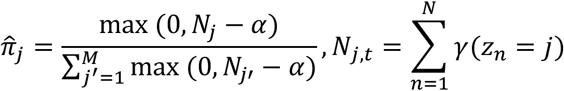

The α parameter can thus be interpreted as the minimum number of ChIP-exo tags required to support a crosslinking event active in the model. MAP values of *µ*_*j*_ are determined by enumerating over several possible values of *µ*_*j*_. Specifically, the MAP estimation of *µ*_*j*_ is:

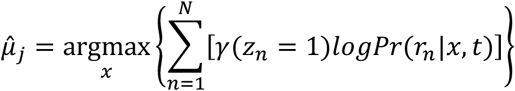

where *x* starts at the previous values of the position weighted by **τ** and expands outwards to 50bp each side. If the maximization step results in two components sharing the same positions, they are combined in the next iteration of the algorithm.

Crosslinking positions are simultaneously modeled across all analyzed ChIP-exo experiments in a given aligned profile, and information about crosslinking position estimates are shared between experiments at each EM step using a positional prior strategy described in the MultiGPS algorithm (Mahony *et al.*, 2014). This positional prior encourages consistency in estimated crosslinking positions across ChIP-exo experiments. Our rationale is that a given protein complex will be crosslinked to DNA at relatively few positions, but the signatures of these crosslinks will be present across experiments characterizing multiple members of the complex.

Finally, the relative crosslinking strengths of all estimated crosslinking points are quantified across all ChIP-exo profiles using Maximum Likelihood assignment of tag counts, yielding a matrix of crosslinking strengths.

### Visualization of crosslinking relationships

The matrix of crosslinking strengths is used to assess the relationships between the crosslinking preferences of each protein in a protein-DNA complex. We normalize the crosslinking strength matrix such that the sum of crosslinking strengths for each protein is 1. We then apply a standard PCA to visualize the relationships between protein-DNA crosslinking preferences.

### Motif analysis

We ran MEME-ChIP version 4.10.0 (Machanick and Bailey, 2011) on tRNA gene sequences to characterize the Box A and Box B motifs. Then, we scan 400bp of regions used for alignment with the discovered motifs using a log-likelihood scoring threshold of 0.1% per base FDR defined using a second-order Markov model based on yeast genome nucleotide frequencies. We obtained the Rap1 cognate DNA-binding motif positions by scanning the corresponding cis-bp database motif (M4379_1.02) (Weirauch *et al.*, 2014) in 1400bp regions centered around TSSs using a log-likelihood scoring threshold of 0.1% per base FDR defined using a second-order Markov model based on yeast genome nucleotide frequencies.

### ChExMix peak calling

We run ChExMix version 0.42 (Yamada *et al.*, 2019) with default parameters on Pol II ChIP-exo data in Kc167 cells and obtained the top 500 most enriched Pol II peaks using q-value.

### Public datasets

ChIP-exo for RPG-specific factors, Tbp1 ChIP-exo, and ChIP-nexus data targeting Pol II and basal TFs are obtained from NCBI Sequence Read Archive under accession number SRP041518 (Reja *et al.*, 2015), GSM2601059 (Vinayachandran *et al.*, 2018), and GSE85741 (Shao and Zeitlinger, 2017), respectively. RPG and Tbp1 ChIP-exo data are aligned against sacCer3 using BWA (Li and Durbin, 2009) version 0.5.9. ChIP-nexus data are aligned against dm3 using BWA version 0.7.12 with options “mem -T 30 -h5”.

### ChIP-exo experiments and processing

The *Saccharomyces cerevisiae* strain, BY4741, was obtained from Open Biosystems. Cells were grown in yeast peptone dextrose (YPD) media at 25°C to an OD_600_=0.8−1.0. ChIP-exo assays were performed as previously described (Rhee and Pugh, 2011; Rossi *et al.*, 2018). Mock IP control ChIP-exo experiments in yeast were performed using rabbit IgG (Sigma, i5006) in the BY4741 background strain (which does not contain a tandem affinity purification tag sequence).

Libraries were paired-end sequenced and read pairs were mapped to the sacCer3 genome using BWA version 0.7.12 with options “mem -T 30 -h 5”. Read pairs that share identical mapping coordinates on both ends are likely to represent PCR duplicates, and so Picard (http://broadinstitute.github.io/picard) was used to de-duplicate such pairs. Reads with MAPQ score less than 5 are filtered out using samtools (Li *et al.*, 2009).

## Availability

Open source code (MIT license) and documentation for ChExAlign are available from https://github.com/seqcode/chexalign. All ChIP-exo sequencing data produced in this study has been uploaded to GEO under accession GSE140923.

### Acknowledgements

This manuscript is based upon work supported by National Institutes of Health grant GM125722 (to S.M. and B.F.P.). Early development of the ChExAlign platform was also supported by the National Science Foundation ABI Innovation Grant No. DBI1564466 (to S.M.). Any opinions, findings and conclusions or recommendations expressed in this material are those of the authors and do not necessarily reflect the views of the National Science Foundation.

